# A Numerical Alternative to MR Thermometry for Safety Validation of Multi-Channel RF Transmit Coils

**DOI:** 10.1101/2025.06.07.657475

**Authors:** Alireza Sadeghi-Tarakameh, Simon Schmidt, Matt Waks, Russell L. Lagore, Andrea Grant, Edward Auerbach, Lance DelaBarre, Lasse Knudsen, Christophe Lenglet, Luca Vizioli, Essa Yacoub, Gregor Adriany, Gregory J. Metzger, Kamil Ugurbil, Yigitcan Eryaman

**Author notes:** Corresponding Author: Alireza Sadeghi-Tarakameh Center for Magnetic Resonance Research (CMRR) University of Minnesota 2021 6th Street Southeast, Minneapolis, MN 55455.

## Abstract

**Purpose:** This study proposes an alternative approach to MR thermometry (MRT) for the safety validation of multi-channel RF transmit coils and demonstrates its use to enable human studies at 10.5T.

**Methods:** To ensure patient safety, specific absorption rate (SAR) limits established under international guidelines must not be exceeded. Predicting SAR on state-of-the-art parallel transmit systems relies on electromagnetic simulations, which require extensive experimental validation. Despite a well-established validation workflow, SAR prediction errors are unavoidable and must be quantified as a safety margin. While MRT tests are commonly used for this purpose, their technical challenges necessitate an alternative. The proposed technique propagates the error between experimentally and numerically acquired *B*^+^distributions to the uncertainty in simulated peak local SAR using Monte-Carlo simulations without the need for MRT. This method was validated using a 16-channel transceiver coil for imaging the human torso (henceforth referred to as a “body” coil) at 10.5T with two excitation scenarios, as well as an 8-channel 10.5T head coil with four excitation scenarios.

**Results:** The proposed numerical technique proved more conservative than existing MRT-based SAR error quantification methods across all tested scenarios. Its application to validate three state-of-the-art head coils (16Tx/32Rx, 16Tx/80Rx, and 16Tx/128Rx) led to regulatory approval for human head imaging and high-quality functional as well as diffusion MRI results at 10.5T.

**Conclusion:** A numerical alternative to MRT requires only the experimental acquisition of *B*^+^ maps for comparison with simulations, enabling the quantification of uncertainty in SAR prediction. This technique was applied to three 16-channel transmit arrays, each used in conjunction with high-channel-count receive arrays for in vivo imaging.

## 1 INTRODUCTION

The increasing interest in high and ultrahigh field (UHF, defined as ≥ 7 Tesla) MRI scanners is primarily driven by their ability to provide a higher signal-to-noise ratio (SNR) at stronger magnetic field strengths,^1–7^ a benefit that is challenging to achieve through other means. A higher SNR enables improved spatial and/or temporal resolution, which is crucial for various clinical and research applications, including anatomical imaging^8–14^ and functional MRI studies (e.g. reviews^15–17^).

Despite the SNR benefits of higher field strengths, several challenges must be overcome to fully exploit their potential. These include excitation inhomogeneity, insufficient radiofrequency (RF) excitation (*B*^+^), and high specific absorption rate (SAR), all of which are primarily linked to RF transmit coils. These issues stem from the higher Larmor frequency associated with stronger main magnetic fields (*B*_0_), which results in shorter wavelengths.

Parallel transmission (pTx) techniques (e.g.^18–24^), which utilize multi-channel RF coils, offer an effective solution by allowing for the channel-wise optimization of *B*^+^fields. Although pTx coils have been shown to be effective across various UHF MRI applications,^14, 24, 25^ commercially available options remain limited. Consequently, researchers often develop custom pTx coils tailored to specific applications(e.g.^26–31^). However, before these coils can be used in human studies, they must undergo stringent patient safety assessments.

According to the latest international guidelines,^32^ which outline the patient safety requirements for RF coils, two parameters must be continuously monitored and controlled during a human scan: global and peak local specific absorption rates. Global SAR is defined as the total RF power dissipated per unit mass of the exposed part of the body, which can be conservatively estimated by monitoring the total input power to the coil. Because MRI scanners are equipped with power monitoring systems, global SAR can be measured in real-time. In contrast, there is currently no reliable method for in-bore measurement of peak local SAR (*pSAR*), making *pSAR* monitoring heavily dependent on electromagnetic (EM) simulations. Since patient safety hinges on the accuracy of these simulations, rigorous experimental validation of pTx coil models is essential. However, despite the clear workflow of modeling and validation^33–35^—which includes reconstructing the 3D model of the pTx coil and comparing simulated versus measured scattering (S) parameters, *B*^+^maps, and local SAR maps in a phantom—errors in modeling and simulations are inevitable. To ensure patient safety, the sources of these errors must be identified, and a safety margin accounting for them should be quantified.

In the literature,^36, 37^ three sources of uncertainty have been identified as contributing to errors in predicting *pSAR* in the human body using EM simulations: power monitoring uncertainty (*e*_*pm*_), inter-subject variability (*e*_*ISV*_),^38–42^ and EM modeling uncertainty (*e*_*EMM*_). Steensma et al.^36^ demonstrated that these uncertainties can be considered uncorrelated and, therefore, can be combined using a sum-of-squares method to calculate the error in predicting *pSAR*, *e*_*SAR*_, as follows:

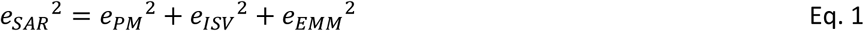

Ultimately, a safety factor (SF, >1) can be calculated using the *e*_*SAR*_ and should be applied to scale the predicted *pSAR* to ensure patient safety, despite the uncertainties:

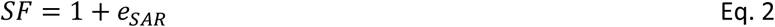

Among the three aforementioned uncertainties, *e*_*pm*_ is often reported by manufacturers, and *e*_*ISV*_ can be determined either by numerical simulations or from literature.^38, 40^ However, to quantify *e*_*EMM*_, the discrepancies between numerically simulated and experimentally measured *B*^+^ and local SAR maps in a tissue-mimicking phantom must be assessed.^33, 34^ Unlike *B*^+^mapping techniques using MR scanners, there is no direct method for local SAR mapping within the scanner. Instead, it can be indirectly calculated using MR thermometry (MRT),^43^ which produces maps of increasing temperature inside the phantom caused by RF energy deposition. The main hurdles of this approach are related to the technical challenges of conducting MRT experiments including the need to perform several different excitation patterns, phase drift errors, system instability, specialized phantoms, and the need for high transmit-power to induce temperature changes— which can be harmful to sensitive electronics, especially in the presence of a dedicated receiver array inside the transmitter.

This study introduces a novel method for quantifying uncertainty in EM modeling, specifically for peak 10g-averaged SAR (*pSAR*_10*g*_) evaluation, without requiring MRT experiments.^44, 45^ Our approach propagates the quantitative differences between measured and simulated *B*^+^ maps to estimate an upper bound for *pSAR*_10*g*_ error. In essence, based on the relationship between the B- and E-fields governed by Maxwell’s equations, we hypothesize that errors in simulated *B*^+^ fields can be used to define the *pSAR*_10*g*_ error space for each excitation mode. We assess the performance of our method against a 16-channel body array and an 8-channel head array validation, which included MRT and temperature probe measurements, respectively.^34, 46^ Additionally, we demonstrate the application of this approach in the safety validation of three high-channel-count head coils at 10.5T and present recent in vivo human brain functional as well as first diffusion imaging data obtained using these coils.

## 2 METHODS

The 10.5T MRI scanner at the Center for Magnetic Resonance Research (CMRR) with a bore diameter of 88cm is the highest field whole body MRI scanner in North America and operates under an Investigational Device Exemption (IDE) from the Food and Drug Administration (FDA). It features a whole-body magnet (Agilent Technologies, Oxford, UK) and an associated imaging system (Siemens Healthineers, Erlangen, Germany), equipped with 16 parallel transmit channels where each is channel driven by a 2-kW RF power amplifier (Stolberg HF-Technik AG, Stolberg, Germany). All RF coils intended for human use in this system require FDA approval, which necessitates rigorous validation of their EM simulation models through experimental measurements.

While the primary focus of this manuscript is a novel approach for quantifying EM modeling uncertainty without relying on MRT, it is important to recognize that this represents only one component of a broader coil validation process. The complete validation workflow for high-channel-count head arrays—which can be generalized to any multi-channel transmit coil—is described first. Next, the proposed EM modeling uncertainty calculation approach is introduced and its performance is evaluated using two RF coils that were previously validated with MRT. Finally, the proposed approach’s integration into the full safety validation pipeline for three high-channel-count neuroimaging arrays at 10.5T is presented.

### 2.1 Safety Validation of Multi-Channel RF Coils

#### 2.1.1 Validation Workflow

The workflow of the safety validation of custom-built multi-channel RF coils can be detailed as:

***1. Scattering (S)-Parameter Measurements:*** Measure the full S-matrix on the bench at the target frequency using a phantom mimicking the dimensions and electrical properties of the targeted body part including the loss and phase shift of each feed cable connected to the coil ports (also those of the T/R switches if used).
***2. EM Simulation and Circuit Model Integration:*** Model the coil in an EM simulation environment (e.g., HFSS, CST, or Sim4life) with its *N*_*l*_ lumped components (e.g., capacitors, inductors) and *N*_*c*_ feed ports, all of which are designated as excitation ports to perform co-simulation.^47^ After exporting the resulting S-matrix (including *N*_*c*_ *+ N*_*l*_ ports) to a circuit simulator (e.g., AWR or ADS), replace each of the *N*_*l*_ lumped component-presenting ports with reactive elements matching the nominal values used in the actual coil. Connect the *N*_*c*_ ports with their respective attenuators (matching measured feed cable losses) and phase shifters (matching measured feed cable phase shifts).
***3. Optimization of Passive Components:*** Perform global and local optimizations to minimize the difference between the measured and simulated *N*_*c*_ -port complex S-matrices. Once satisfactory agreement is achieved between the measured and simulated S-parameters, replace the circuit simulator ports with voltage sources featuring a 50Ω internal resistance. Activating one-by-one using a 1V-excitation, the resulting voltages across all *N*_*c*_ attenuators were recorded as an *N*_*c*_ × 1 vector per enabled voltage source resulting in *N*_*c*_ excitation vectors.
***4. Per-Channel Excitations in EM Simulation:*** In the EM simulation software, terminate the *N*_*l*_ ports with reactance values matching the optimized lumped components from the previous step. Generate the per-channel excitation corresponding to the *i*^*th*^ channel of the actual coil, use the *i*^*th*^ excitation vector (*N*_*c*_ × 1) obtained from the circuit simulator to excite the *N*_*c*_ ports of the simulation model. Export the per-channel complex *B*^+^ and 𝐸-field distributions along with the mass density and conductivity distributions of the phantom.
***5. Q-Matrix Calculation:*** Calculate the10g-averaged local SAR matrices (𝑄-matrices) using the exported *E*-fields, mass density, and conductivity data.^48, 49^
***6.*** 𝑩_1_^+^ ***Measurements:*** Experimentally measure the relative per-channel complex *B*^+^ distributions, as well as the absolute *B*^+^ map of a selected excitation mode, such as the circularly-polarized (CP) mode, in a physical phantom used for simulation.
***7. Safety Factor Calculation:*** Using our new proposed approach (outlined in Section 2.1.2 and Figure 1) estimate the EM modeling uncertainty (*e*_*EMM*_) using the simulated and measured *B*^+^ maps and simulated 𝑄-matrices As previously suggested,^36^ inter-subject variability (*e*_*ISV*_) can be determined using numerical techniques^38^ or selected from relevant literature (e.g., Refs^40, 41^). Power monitoring uncertainty (*e*_*pm*_) values, typically reported by the vendor, range from 10% to 15%. These three uncertainty values (*e*_*pm*_, *e*_*ISV*_, *e*_*EMM*_) are combined in a sum-of-squares fashion to calculate the safety factor (𝑆𝐹), as formulated in Equations 1 and 2.
***8. Human Model Simulation:*** Repeat Steps 4-5 (the per-channel *B*^+^/*E*-field and 𝑄-matrix generation steps) in a human body model (e.g., Duke, Ella, or ANSYS Human Model) allowing for the creation of Virtual Observation Points (VOPs)^48^ scaled using the 𝑆𝐹 calculated in Step 7 to account for different sources of uncertainty and ensure compliance with safety guidelines.

**Figure 1.**
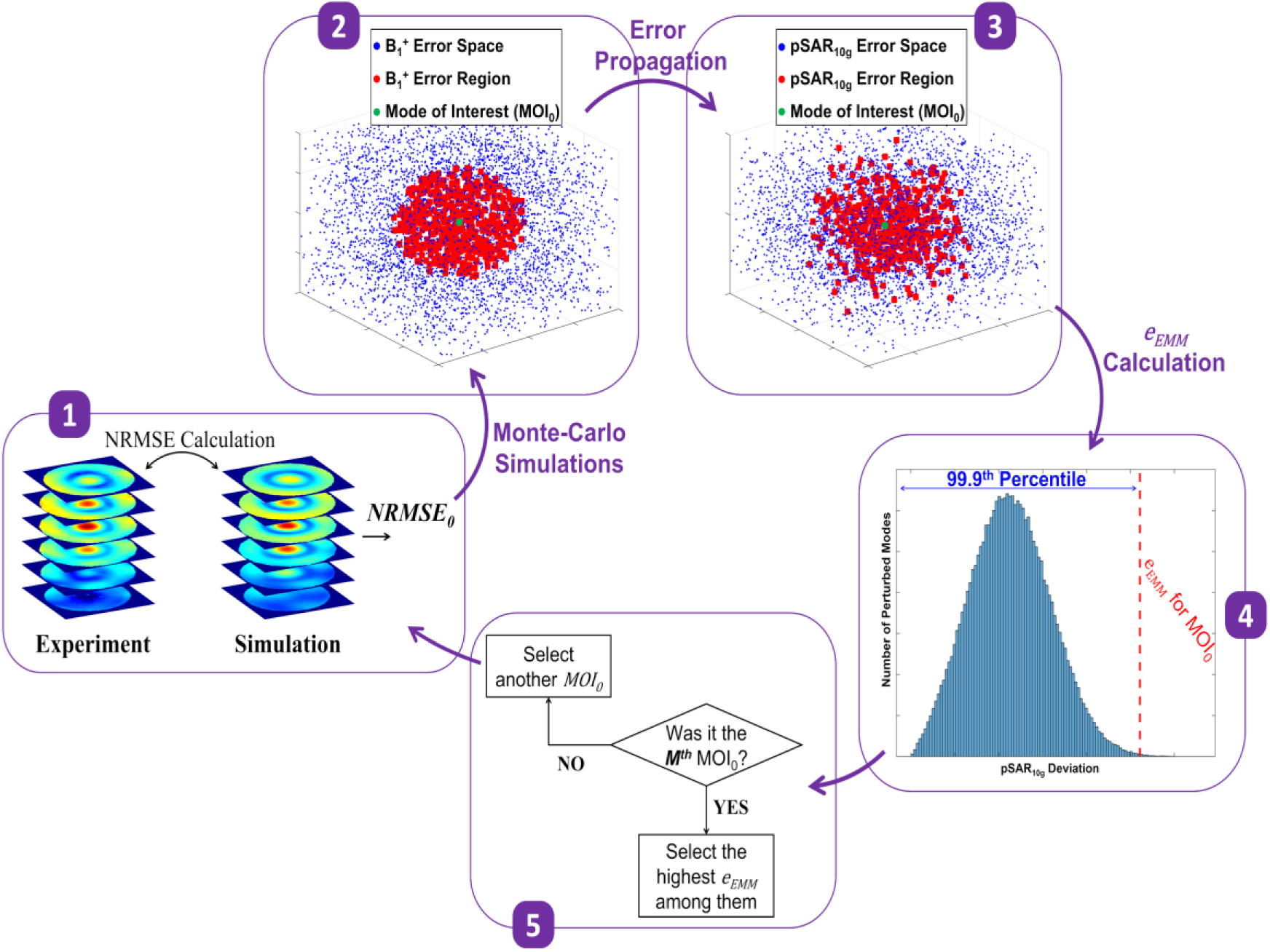
Mockup illustration of the proposed five-step approach for quantifying EM modeling uncertainty in predicting *pSAR*10*g* generated by a multi-channel RF transmit coil, based on the error between simulated and measured *B*+ maps. (A) Simulated and measured *B*+ maps are generated for a mode of interest using per-channel *B*+ data, and the error between them is computed as NRMSE0. (B) Monte Carlo simulations are performed by perturbing the magnitude and phase of each channel around the mode of interest, generating the *B*+ error space. Modes satisfying NRMSEi ≤ NRMSE0 are selected to form the *B*+ error region around the mode of interest. (C) The *B*+ error space and region are propagated to generate corresponding *pSAR*10*g* error space and region. (D) The *pSAR*10*g* error region is analyzed, and the 99.9th percentile of the distribution is identified as the as the *e*_*EMM*_ for the selected mode of interest. (E) The entire process is repeated for additional modes of interest to evaluate global modeling uncertainty.

#### 2.1.2 EM Modeling Uncertainty Calculation

Figure 1 presents a mockup illustration summarizing the five-step approach proposed in this work for quantifying EM modeling uncertainty (Step 7 above). This method numerically propagates the error between simulated and measured *B*^+^ fields to estimate the uncertainty in predicting the real-life *pSAR*_10*g*_ value. The steps are outlined as follows:

1. Generate a Mode of Interest (MOI0)

***a.*** Using a desired set of magnitudes and phases, generate the corresponding 3D *B*^+^ maps for both simulated and experimental per-channel *B*^+^ data.
***b.*** Calculate the normalized root-mean-square error (*N*𝑅𝑀𝑆*E*_0_) between the 3D *B*^+^ maps obtained from the experimental and simulated data.
2. Perform Monte-Carlo Simulations (N iterations; i=1:N)

***a.*** In the *i*^*th*^ iteration, perturb the complex excitation vector in the vicinity of 𝑀𝑂𝐼_0_ to generate the perturbed mode 𝑀𝑂𝐼_*i*_.
***b.*** Compute the NRMSEᵢ between the 3D *B*^+^ maps of 𝑀𝑂𝐼_*i*_ (perturbed mode) and 𝑀𝑂𝐼_0_ (unperturbed mode) to construct 𝑀𝑂𝐼_0_’s *B*^+^ error space.
***c.*** Select all perturbed modes (𝑀𝑂𝐼_*i*_) satisfying *N*𝑅𝑀𝑆*E*_i_ ≤ *N*𝑅𝑀𝑆*E*_0_ and generate the *B*^+^ error region.
3. *Propagate* 𝑩_1_^+^ *Error Space to* 𝒑𝑺𝑨𝑹_𝟏𝟎𝒈_ *Error Space*

***a.*** Compute the *pSAR*^*i*^ for all perturbed modes (𝑀𝑂𝐼_*i*_) identified in Step 2a.
***b.*** Calculate the error between *pSAR*^*i*^ values and *pSAR*^0^ of 𝑀𝑂𝐼_0_, generating the 𝑀𝑂𝐼_0_’s *pSAR*_10*g*_ error space.
***c.*** Identify *pSAR*^*i*^ values corresponding to 𝑀𝑂𝐼_*i*_ modes satisfying Step 2c and generate the *pSAR*_10*g*_ error region.
4. *Determine the 99.9th Percentile of the* 𝒑𝑺𝑨𝑹_𝟏𝟎𝒈_ *Error Region as the* 𝒆_𝑬𝑴𝑴_ *of* 𝑴𝑶𝑰_𝟎_

***a.*** Generate a histogram of the *pSAR*_10*g*_ uncertainty region from Step 3c.
***b.*** Select the 99.9th percentile of the histogram as the *e*_*EMM*_ of 𝑀𝑂𝐼_0_.
5. *Repeat Steps 1–4 for* 𝑴 *Different* 𝑴𝑶𝑰_𝟎_ *and Select the Highest* 𝒆_𝑬𝑴𝑴_

Note that the five-step approach outlined above provides a general framework for *e*_*EMM*_ quantification. The technical details of its implementation, as applied to the safety validation of multiple transmit array coils at 10.5T, are discussed in the subsequent sections.

### 2.2 Performance Evaluation of the Proposed *e*_*EMM*_ Calculation Approach

#### 2.2.1 Reference to the MRT Technique

To evaluate the efficacy of the proposed approach in quantifying EM modeling uncertainty for predicting *pSAR*_10*g*_, two modes of interest (𝑀𝑂𝐼_0_)—CP and alternating-phase (*0°-180°-0°-…*)—were investigated using a 16-channel dipole transceiver body coil^46^ at 10.5T (Figure 2A). The study was conducted with a uniform torso-sized phantom (CLP191 Phantom, The Phantom Laboratory Incorporated, Greenwich, CT), characterized by σ = 0.61 S/m and εr = 45.7.

**Figure 2.**
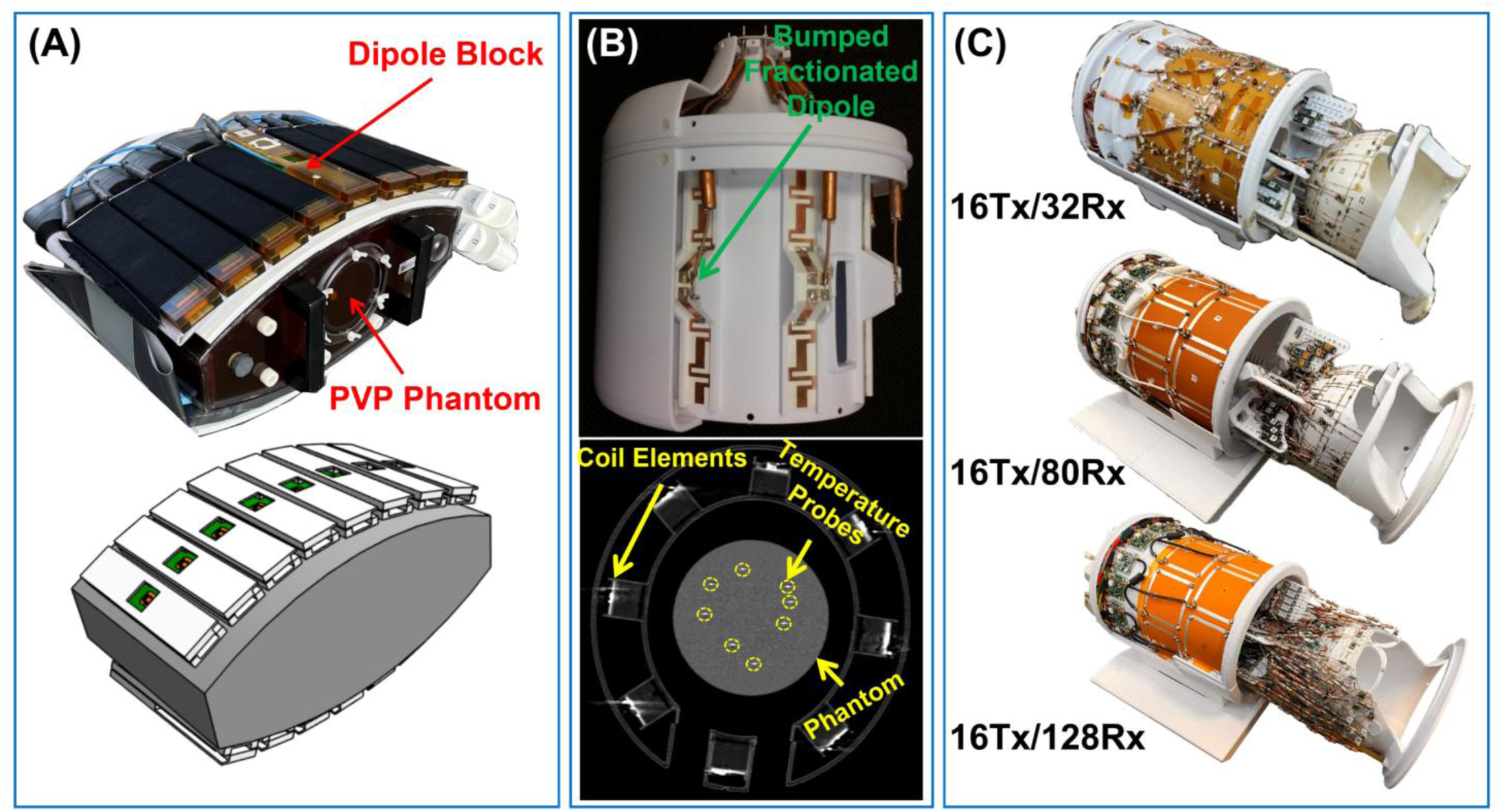
Multi-channel RF transmit coils designed for 10.5T, used either to evaluate the proposed *e*_*EMM*_ calculation technique or to undergo safety validation using the technique. (A) Experimental setup and simulation model of the 16TxRx body coil, previously safety-validated using *B*+ mapping and MRT tests, and used in this study as a reference to evaluate the efficacy of the proposed *e*_*EMM*_ calculation. (B) Experimental setup and CT image of an 8TxRx head coil, previously safety-validated using *B*+ mapping and temperature probe measurements, and used in this study to evaluate the proposed technique’s performance across different random modes of interest. Temperature probe locations are shown on the CT image. (C) Three high-channel-count head coils (16Tx/32Rx, 16Tx/80Rx, and 16Tx/128Rx) that were safety-validated using the proposed technique and used for in vivo functional and diffusion MRI at 10.5T.

The *e*_*EMM*_s computed using the proposed method were compared to those obtained via the MRT-based technique.^36^ Notably, in the MRT-based technique, voxel-wise comparisons between simulated and experimental *B*^+^ and SAR maps were used for *e*_*EMM*_ quantification. In contrast, our approach propagates the *N*𝑅𝑀𝑆*E* between measured and simulated *B*^+^ maps of a given excitation mode into the *pSAR*_10*g*_ error space of that mode.

Error propagation was performed using 10⁵ Monte Carlo simulations, following Steps 2 and 3 in Figure 1. The magnitude of per-channel *B*^+^ perturbations was constrained within +/- the per-channel *B*^+^ errors — *NRMSE* between per-channel simulated and measured *B*^+^maps. These 10⁵ perturbed modes in the vicinity of the chosen MOI were then used to construct the *pSAR*_10*g*_ error region. Finally, the 99.9th percentile of the *pSAR*_10*g*_ error region was identified as the *e*_*EMM*_ of that mode (Step 4 in Figure 1). The *e*_*EMM*_s calculated via this numerical approach were subsequently compared to those obtained from experimentally measured SAR in Schmidt et al.’s work.^46^

The per-channel *B*^+^ maps were acquired using a hybrid technique,^50, 51^ combining:

- Relative *B*^+^ mapping^50^: 2D axial GRE with TE/TR = 2.48/50 ms, FA = 7°, FOV = 470 × 234 × 5 mm^3^, Resolution = 1 × 1 × 5 mm^3^, and pixel bandwidth = 970 Hz/px.
- Absolute *B*^+^ mapping (Actual Flip-Angle Imaging)^52^: TE/TR1/TR2 = 1.72/28/128 ms, FA = 90°, FOV = 470 × 234 × 240 mm^3^, Resolution = 3 × 3 × 5 mm^3^, and pixel bandwidth = 1110 Hz/px.

MRT tests were conducted using 10 minutes of RF-induced heating at 88 W time-averaged power, followed by two multi-echo GRE acquisitions (TE = 5/10/15 ms) recorded before and after RF exposure. SAR was then computed from the temperature change (Δ𝑇) using the formula:

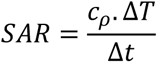

where *c*_𝜌_ denotes the specific heat capacity (measured as 3500 J/kg/K) and Δ𝑡 =600s is the RF exposure duration.

#### 2.2.2 Reference to the Temperature Probing Technique

To further validate the efficacy of the proposed technique—particularly for excitation modes beyond those used for

*e*_*EMM*_ calculations—an eight-channel transceiver (TxRx) 10.5T head coil^34^ was tested using a uniform jar-shaped phantom (σ = 0.5 S/m, εr = 78) equipped with eight temperature probes (Figure 2B). For this validation, the CP mode of interest (𝑀𝑂𝐼_0_) was first utilized, and its corresponding *e*_*EMM*_ was calculated (Steps 1–4 in Figure 1), targeting 10g-averaged SAR at the eight probe locations. In other words, the *Q*-matrix was reduced to eight discrete points, and the resulting *e*_*EMM*_ merely reflected the modeling uncertainty in predicting *pSAR*_10*g*_ at those specific locations. This process was then repeated for three additional excitation modes: linear excitation, zero-phase excitation, and a random excitation. Ultimately, the highest *e*_*EMM*_ among the four tested modes was selected as the final *e*_*EMM*_ for this coil model (Step 5 in Figure 1).

To assess the accuracy of the calculated *e*_*EMM*_, the coil was driven with three additional random excitations, and the resulting temperature increase inside the phantom was measured at eight arbitrary locations (indicated with yellow circles on Figure 2B). The same excitation modes were then applied in simulations, and the local SAR data was recorded at the same eight locations. Finally, the simulated SAR data were scaled using the calculated *e*_*EMM*_ and compared against the measured SAR values.

All *B*^+^ maps were acquired using the AFI technique.^52^ Temperature measurements were performed using eight fiber optic temperature probes (Lumasense Technologies, CA). The slope of the initial linear region of the temperature progression curve was then used to estimate the local SAR at the probe tips.

### 2.3 Case Studies: Safety Validation of three 10.5T Head Coils

The development of RF technology for head imaging at 10.5T has seen steady and continuous growth over the past decade. This progress evolved from the first 8-channel transceiver head coil^34^ (Figure 2B), initially developed for feasibility studies, to the state-of-the-art 16Tx/80Rx^53^ and 16Tx/128Rx coils.^54^ As mandated by the FDA under an IDE, all developed coils underwent rigorous safety validation procedures before human use. The following provides technical details of the safety validation workflow implementation (see Section 2.1.1) for the three high-channel-count head coils developed for neuroimaging at 10.5T:

<u>EM Simulations:</u> The EM models of three high-channel-count 10.5T head coils that are shown in Figure 2C—16Tx/32Rx,^55, 56^ 16Tx/80Rx,^53^ and 16Tx/128Rx^54^—were developed in an EM simulation environment (HFSS, ANSYS Inc., Canonsburg, PA). The solution type was set to “*HFSS: Network Analysis*” with “*Auto-Open Region: Radiation*” enabled. The excitation ports were defined as “*Terminal Lumped Port*”, and analysis settings were configured based on the coil model’s structural characteristics as:

- *Single frequency of interest*
- *Maximum number of passes:* 20
- *Max. Delta S convergence:* 10^-3^
- *Mixed Order Basis Functions* with *Auto Select Solver*

These simulations were conducted using a uniform lightbulb-shaped phantom with approximate human head and neck-like dimensions (σ = 0.65 S/m, εr = 47.2). The receive-insert arrays were not included in the EM models, as they were demonstrated to have minimal impact on the transmit coils.

<u>Optimization of S-parameters:</u> The S-parameters of the 16-channel transmitters were measured on the bench using a 16- channel network analyzer (Rohde & Schwarz ZNBT8, Munich, Germany) while their respective receive-insert coils were in place. Using a circuit simulator (AWR, Cadence Design Systems Inc., San Jose, CA) and its built-in optimization toolbox, a two-step optimization (i.e., global and local optimization) was conducted to match complex-valued S-parameters of the EM models to the bench-top measurements. The global optimization using the *Advanced Genetic Algorithm* and the local optimization using the *Gradient Descent* technique were performed to minimize the least-squares (LS) difference between the measured and simulated S-matrices.

B_1_^+^ Measurements: For each 16Tx array, the per-channel complex *B*^+^ maps were obtained using the hybrid *B*^+^ mapping technique^51^—a fast relative *B*^+^ mapping^50^ scaled using the absolute *B*^+^ maps acquired by the AFI data^52^—while the receive-insert coils were in-place. These *B*^+^ field maps were then used to generate parallel transmission (pTx) excitation patterns.

<u>Safety Factor Calculation:</u> The *e*_*EMM*_s for the three head coils were calculated using the numerical approach proposed in this work (Figure 1). These values were then combined with a 50% inter-subject variability^40^ (*e*_*ISV*_) and a 15% power monitoring uncertainty^46^ (*e*_*pm*_) to determine the safety factor. To calculate the *e*_*EMM*_, the CP mode was selected as the 𝑀𝑂𝐼_0_, and the *N*𝑅𝑀𝑆*E* between the measured and simulated *B*^+^ maps of this mode was propagated to the *pSAR*_10*g*_ error space. The error propagation was performed using 10⁵ Monte-Carlo simulations (Step 2 in Figure 1), with per-channel perturbations constrained within +/- the per-channel *B*^+^ errors. The resulting perturbed modes (10⁵ simulations) were used to construct the *pSAR*_10*g*_ error region, and the 99.9th percentile of this region was selected as the *e*_*EMM*_ (Step 4 in Figure 1).

<u>Human Model Simulation:</u> A human head model (ANSYS Very-high Precision Male model, ANSYS Inc., Canonsburg, PA) was simulated inside the EM models of the three head coils. The corresponding 𝑄-matrices were computed and compressed into VOPs with a 10% overestimation. Finally, the VOPs for each coil were scaled using their respective safety factors and incorporated into a coil-specific file loaded by the system when the coil is plugged in. The RF safety watchdog (power monitoring system) utilizes these embedded VOPs to calculate a predicted *pSAR* and provide real-time *pSAR* monitoring.

### 2.4 Human Brain In Vivo Functional and Diffusion Imaging at 10.5T

#### 2.4.1 Functional MRI

One of the primary motivations behind the deployment of the 10.5T scanner at CMRR has been to enable the acquisition of high-resolution functional MRI (fMRI) maps of the human brain with enhanced functional sensitivity, leveraging the unprecedented SNR available at this field strength. From the early development of the first safety-validated 8-channel transceiver head coil^57^—initially built for feasibility studies—this goal has been consistently pursued. The desire to harness even greater SNR drove the development of high-channel-count, state-of-the-art head coils, which in turn introduced new safety validation challenges. These challenges were addressed through the novel techniques proposed in this work, ultimately enabling the acquisition of high-quality fMRI data at 10.5T using these safety-validated, high-channel-count coils.

fMRI data with submillimeter resolution were acquired at 10.5T using four safety-validated head coils: 8TxRx, 16Tx/32Rx, 16Tx/80Rx, and 16Tx/128Rx. The respective resolutions were 0.54 × 0.54 × 0.8 mm³, 0.4 × 0.4 × 0.6 mm³, 0.5 mm isotropic, and 0.35 mm isotropic, acquired using 2D and 3D GRE-EPI sequences.

Further details on fMRI data acquisition and processing are available in Supporting Information.

#### 2.4.2 Diffusion MRI

Diffusion MRI (dMRI) at high magnetic fields, while presenting an opportunity for access to higher SNR, can be severely limited by power deposition, transmit field uniformity, and even peak power due to the need for a 180° refocusing pulse. Consequently, the feasibility and benefits of high-field dMRI are tightly coupled to the power and safety limits defined for a given RF coil. To demonstrate the utility of the methods proposed in this work—and despite concerns that the technique’s conservative estimation of *e*_*EMM*_ might result in insufficient *B*^+^—the first in vivo human brain dMRI data at 10.5T were acquired using two safety-validated high-channel-count head coils: 16Tx/80Rx and 16Tx/128Rx. Whole-brain dMRI was performed with 1.05 mm isotropic resolution using a two-shell protocol with *b*-values of 900 and 1800 s/mm².

Further details on dMRI data acquisition and processing are available in Supporting Information.

## 3 RESULTS

### 3.1 Performance Evaluation of the Proposed *e*_*EMM*_ Calculation Approach

#### 3.1.1 Reference to the MRT Technique

Figure 3 summarizes the performance evaluation study of the proposed numerical *e*_*EMM*_ calculation technique, conducted using the CP mode of the 16-channel transceiver body coil shown in Figure 2A. Specifically, Figure 3A-B display the measured and simulated *pSAR*_10*g*_ maps on an axial slice inside the phantom, corresponding to the CP mode of interest. The 99.9th percentile of the voxel-wise deviation between the simulated and measured SAR (Figure 3E) was considered the ground truth *e*_*EMM*_ (36%). In contrast, Figure 3F illustrates the distribution of the numerically predicted deviation of the simulated *pSAR*_10*g*_ from its real-life value in the CP mode of excitation. This distribution was obtained by propagating the 28% *N*𝑅𝑀𝑆*E* (i.e., details of the calculation are available in^46^) between the measured and simulated *B*^+^maps (Figure 3C-D) into the *pSAR*_10*g*_ error region, following the methodology described in Section 2.1.2. Ultimately, the 99.9th percentile of the *pSAR*_10*g*_ error region was selected as the numerically predicted *e*_*EMM*_ (49%).

**Figure 3.**
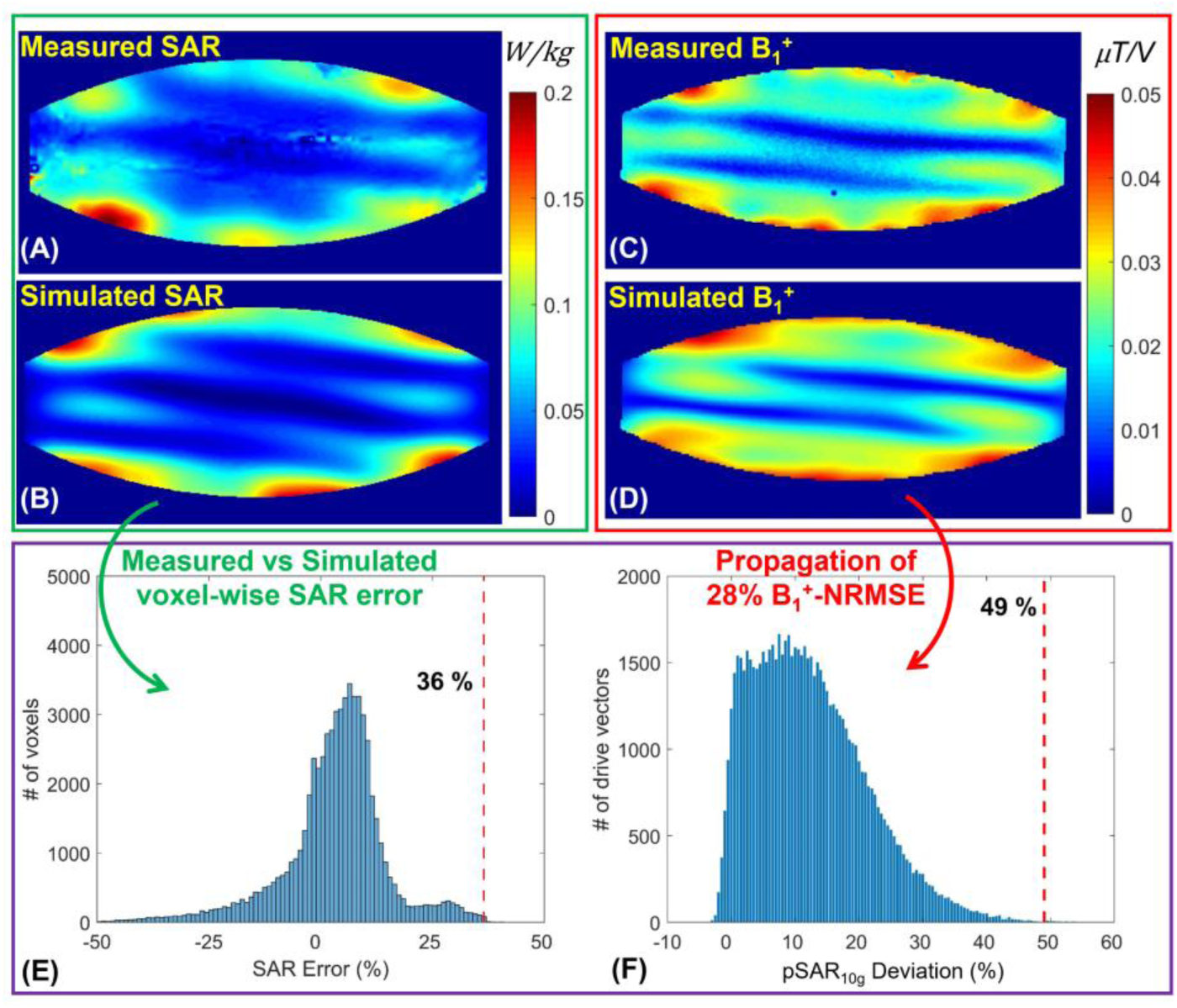
Evaluation of the proposed *e*_*EMM*_ calculation technique using the CP excitation mode of the 16TxRx body coil. (A–B) Measured (via MRT) and simulated local *SAR*10*g* maps. (C–D) Measured and simulated *B*1+ maps for the same excitation mode. *e*_*EMM*_ quantification (E) based on MRT-derived SAR measurements and (F) by propagating the *B*1+ error into the *pSAR*10*g* error space using the proposed method.

The same evaluation was performed for the alternating-phase mode of excitation, with results presented in Figure 4. In this case, the ground truth *e*_*EMM*_, determined via MRT, was 40%, while a 36% *N*𝑅𝑀𝑆*E* between the simulated and measured *B*^+^maps was propagated to a 46% *e*_*EMM*_ using the proposed technique.

**Figure 4.**
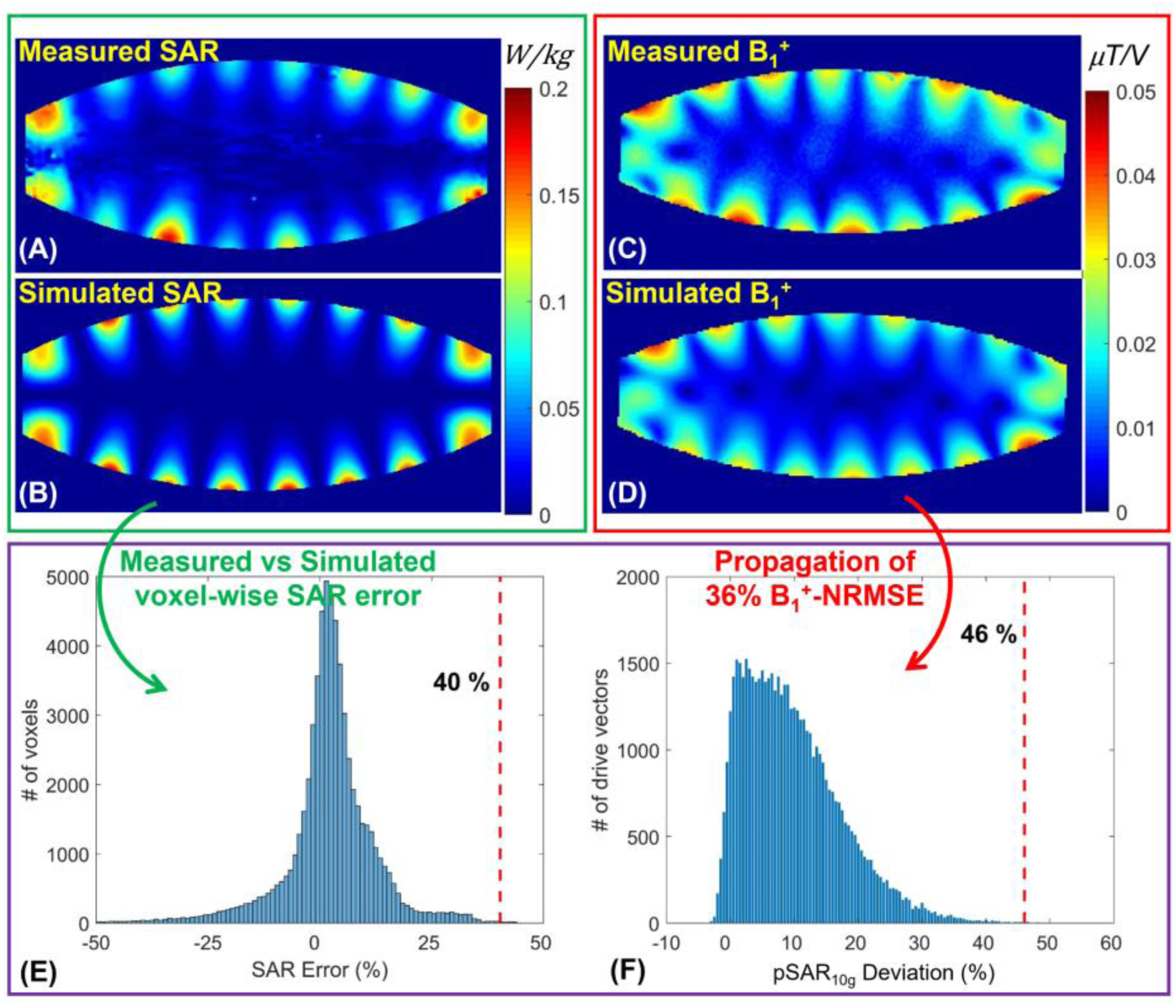
Evaluation of the proposed *e*_*EMM*_ calculation technique using the alternating-phase excitation mode of the 16TxRx body coil. (A–B) Measured (via MRT) and simulated local *SAR*10*g* maps. (C–D) Measured and simulated *B*1+ maps for the same excitation mode. *e*_*EMM*_ quantification (E) based on MRT-derived SAR measurements and (F) by propagating the *B*1+ error into the *pSAR*10*g* error space using the proposed method.

#### 3.1.2 Reference to the Temperature Probing Technique

Figure 5A presents the experimentally measured and numerically simulated *B*^+^ maps for four different excitation modes corresponding to the 8-channel bumped dipole head coil (Figure 2B). The *N*𝑅𝑀𝑆*E*s between the simulated and measured maps were propagated numerically into the corresponding *pSAR*_10*g*_ error regions, as shown in Figure 5B. For each excitation mode, the 99.9th percentile of the *pSAR*_10*g*_ error region is marked on the corresponding histogram as the *e*_*EMM*_ for that mode. Among the four excitation modes, the highest *e*_*EMM*_ was 43%, which was selected as the overall *e*_*EMM*_.

**Figure 5.**
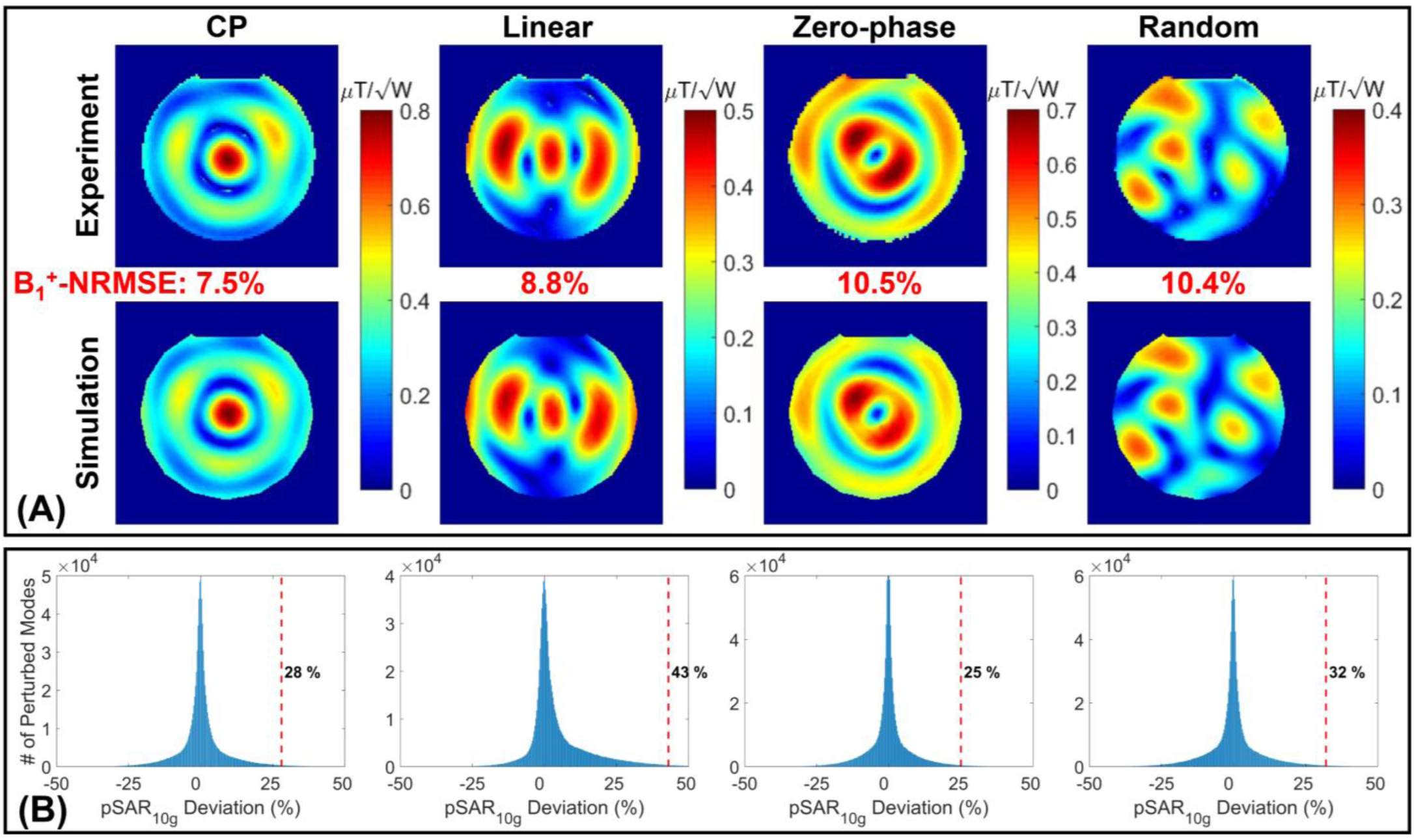
Propagation of the error between measured and simulated *B*1+ maps into the uncertainty in *pSAR*10*g* prediction at the eight temperature probe locations for the 8TxRx head coil using four different excitation modes. (A) Measured and simulated *B*1+ maps corresponding to the four excitation modes. (B) Distributions of the *pSAR*10*g* error region for each mode, as defined in Steps 3 and 4 of Figure 1.

Figure 6A-C displays the simulated *pSAR*_10*g*_ maps for three arbitrary excitation patterns. In Figure 6D-F, the red and blue bars represent the measured *pSAR*_10*g*_ values (obtained using temperature probes) and the simulated *pSAR*_10*g*_ values at the locations of eight temperature probes. The simulated *pSAR*_10*g*_ values were then scaled up by 43%—corresponding to the *e*_*EMM*_—and are shown as green bars in Figure 6D-F.

**Figure 6.**
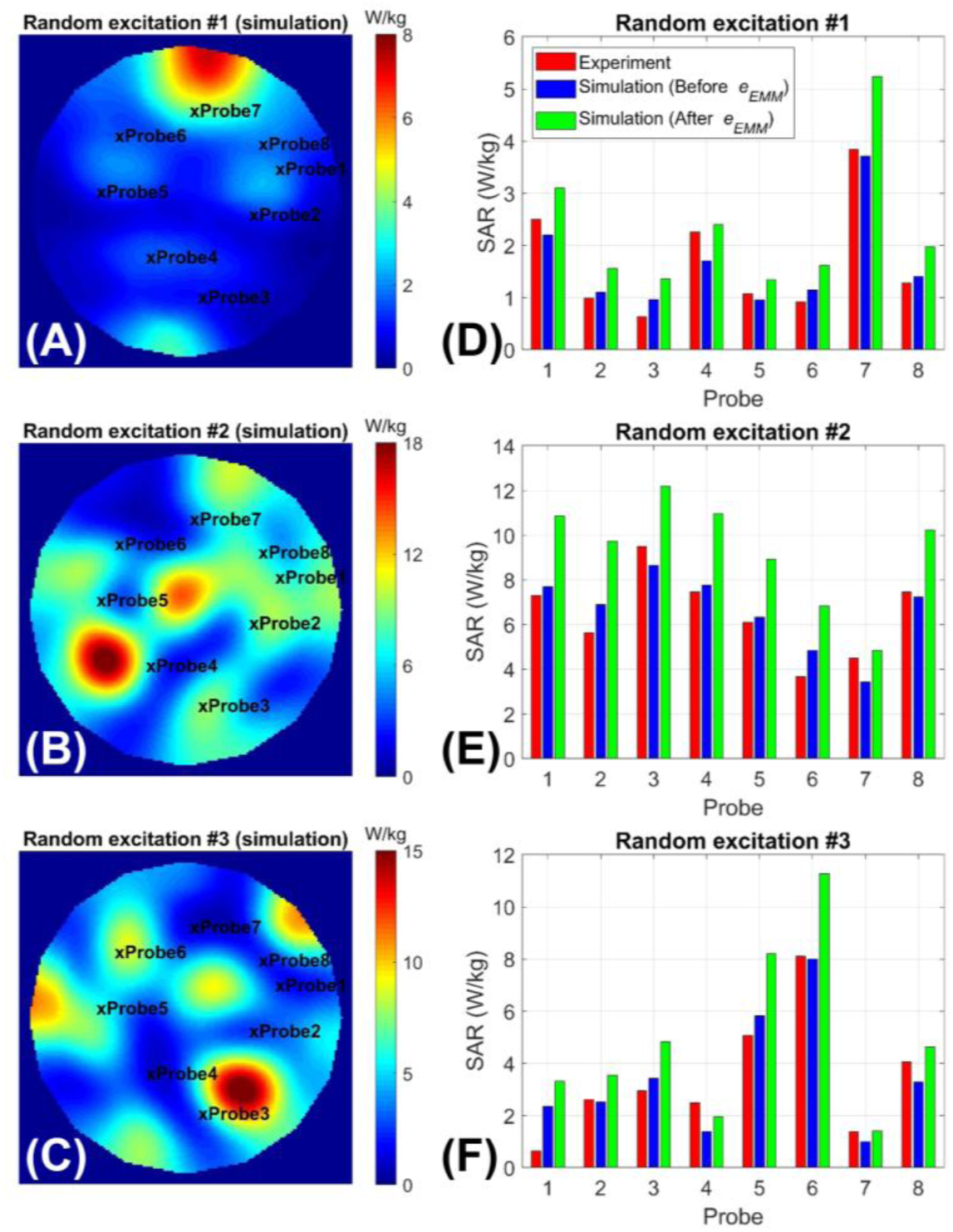
Assessment of the efficacy of the *e*_*EMM*_, calculated using the proposed technique applied to the four excitation modes in Figure 5, and used as a safety margin on simulated SAR₁₀g at the locations of eight temperature probes. (A–C) Simulated SAR₁₀g distributions on an axial plane inside the phantom for three arbitrary excitation modes. (D–F) Comparison of measured (via temperature probe measurements) and simulated *SAR*10*g* values at the eight probe locations. Blue bars represent the simulated *SAR*10*g* before applying the safety margin, while green bars show values after scaling with the 43% *e*_*EMM*_. Notably, no underestimation of *pSAR*10*g* was observed after scaling, confirming the conservativeness of the proposed approach.

### 3.2 Case Studies: Safety Validation of three 10.5T Head Coils

Figure 7A–C presents the measured and simulated S-matrices for three 10.5T transmit array coils shown in Figure 2C— 16Tx/32Rx,^55, 56^ 16Tx/80Rx,^53^ and 16Tx/128Rx.^54^ Measurements were conducted on the benchtop with their respective receive-insert arrays in place, whereas simulations were performed without the receive coils and subsequently co-simulated to match the measured complex S-parameters.

**Figure 7.**
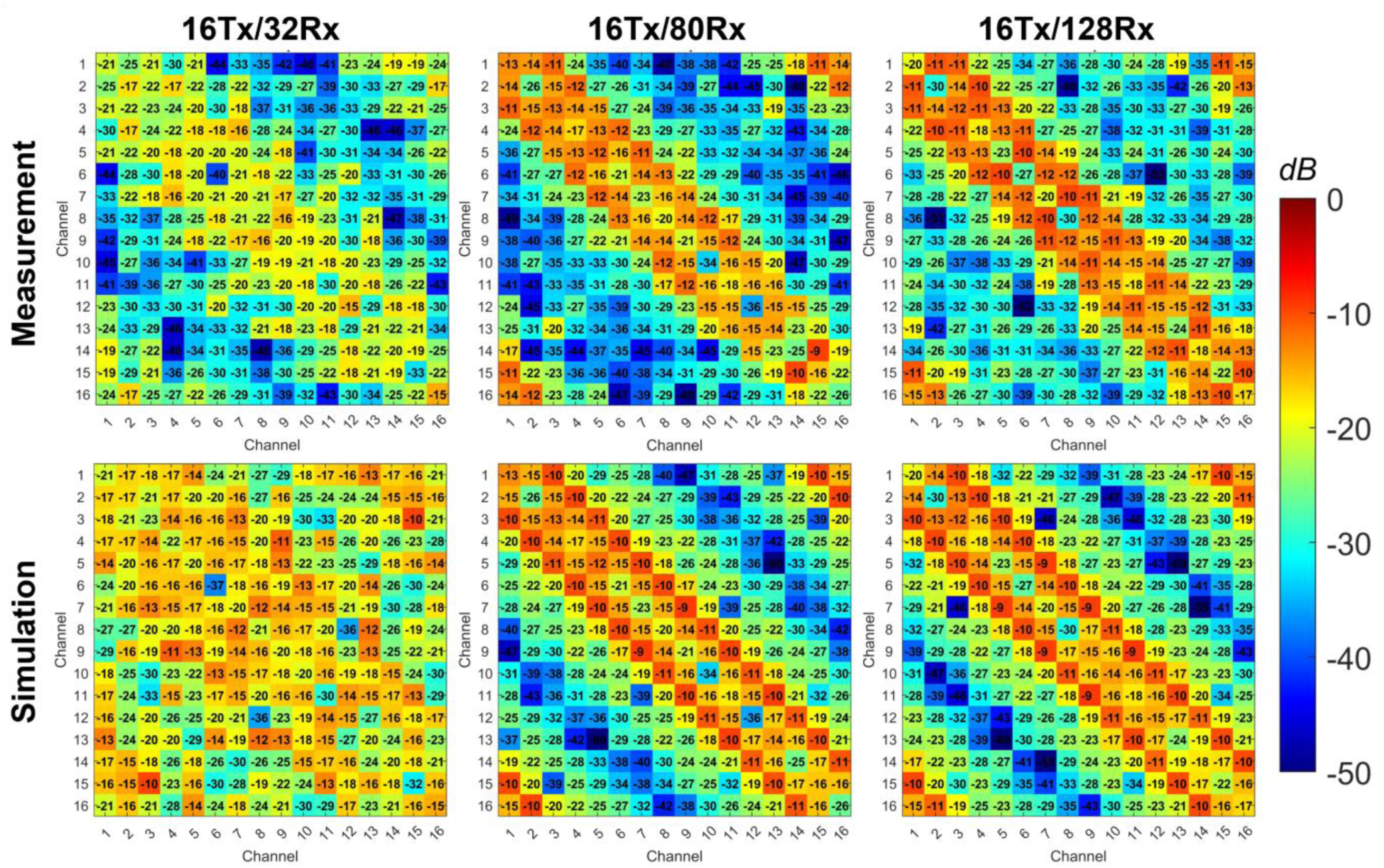
Bench-measured (top row) and simulated (bottom row) S-parameters (in dB) for the 16-channel transmit arrays of the three safety-validated head coils at 10.5T: 16Tx/32Rx, 16Tx/80Rx, and 16Tx/128Rx. All measurements were performed with the coils loaded by the lightbulb-shaped phantom and their respective receive-insert coils in place.

Figure 8 (top row) displays the experimentally measured and numerically simulated *B*^+^ maps for the CP mode of excitation in the three 10.5T head coils. The *N*𝑅𝑀𝑆*E*s between the simulated and measured *B*^+^maps were propagated numerically into the corresponding *pSAR*_10*g*_ error regions, as shown in Figure 8 (bottom row). For each coil, the 99.9th percentile of the *pSAR*_10*g*_ error region is highlighted on the corresponding histogram as the *e*_*EMM*_ for that coil. Using Equations 1 and 2, these *e*_*EMM*_s were combined with a 50% inter-subject variability (*e*_*ISV*_) (from literature^40^) and a 15% power monitoring uncertainty (*e*_*pm*_) (reported by the vendor) to calculate safety factors of 2.42, 1.71, and 1.92 for the three coils, respectively.

**Figure 8.**
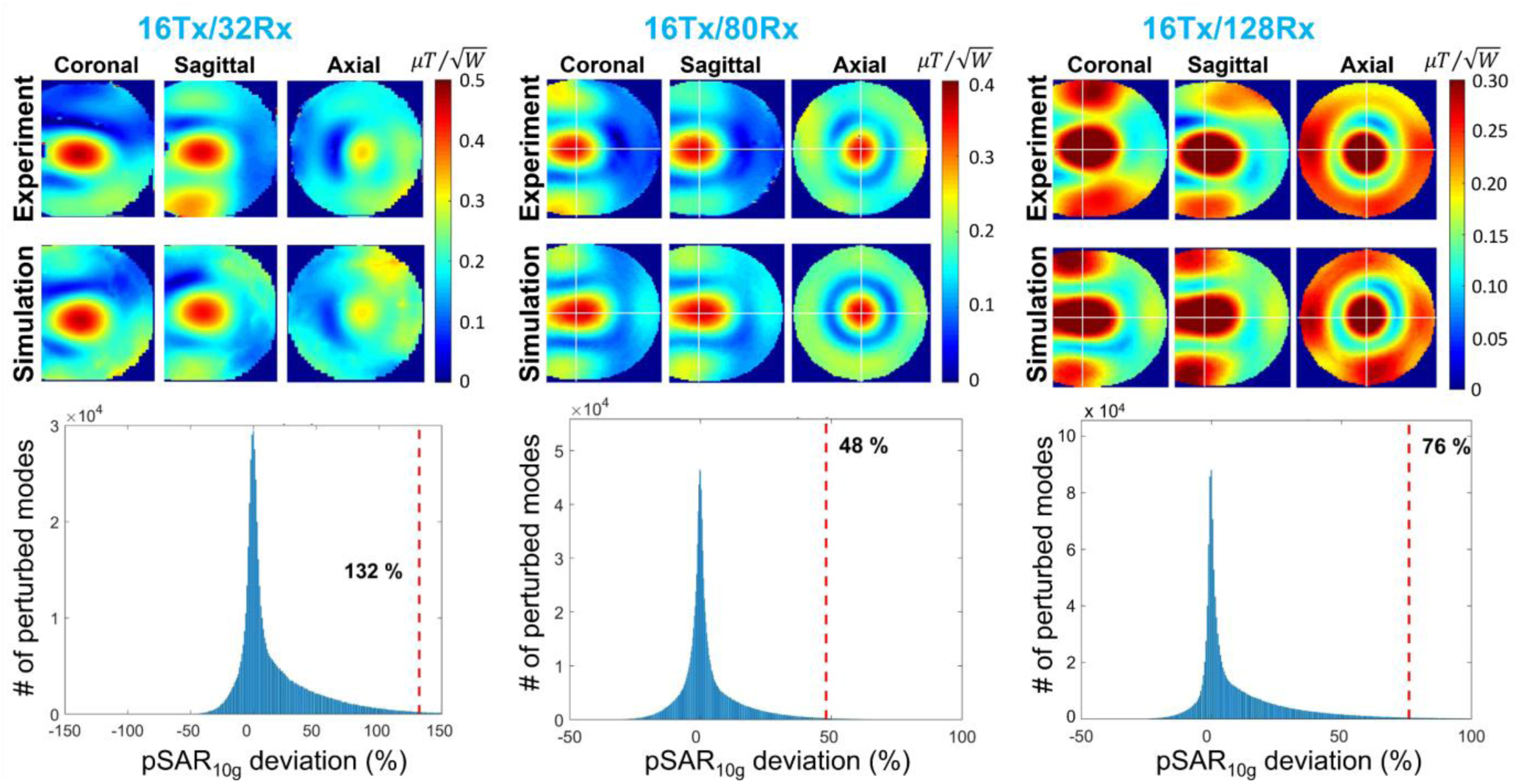
Propagation of the error between measured and simulated *B*1+ maps into the uncertainty in *pSAR*10*g* prediction for three high-channel-count head coils (16Tx/32Rx, 16Tx/80Rx, and 16Tx/128Rx) at 10.5T using their CP mode of excitation. Top row: Measured and simulated *B*1+ maps for each coil. Bottom row: Corresponding distributions of the *pSAR*10*g* error regions, illustrating the resulting *e*_*EMM*_ for each coil.

### 3.3 Human Brain In Vivo Anatomical and Functional Imaging at 10.5T

#### 3.3.1 Functional MRI

Figure 9A–D presents 10.5T fMRI data, showing functional maps acquired during a visual stimulation experiment using four safety-validated head coils. Functional maps with in-plane resolution as fine as 0.4 × 0.4 mm² were obtained using the 2D GRE-EPI sequence, while isotropic resolutions down to 0.35 mm were achieved with the 3D GRE-EPI sequence.

**Figure 9.**
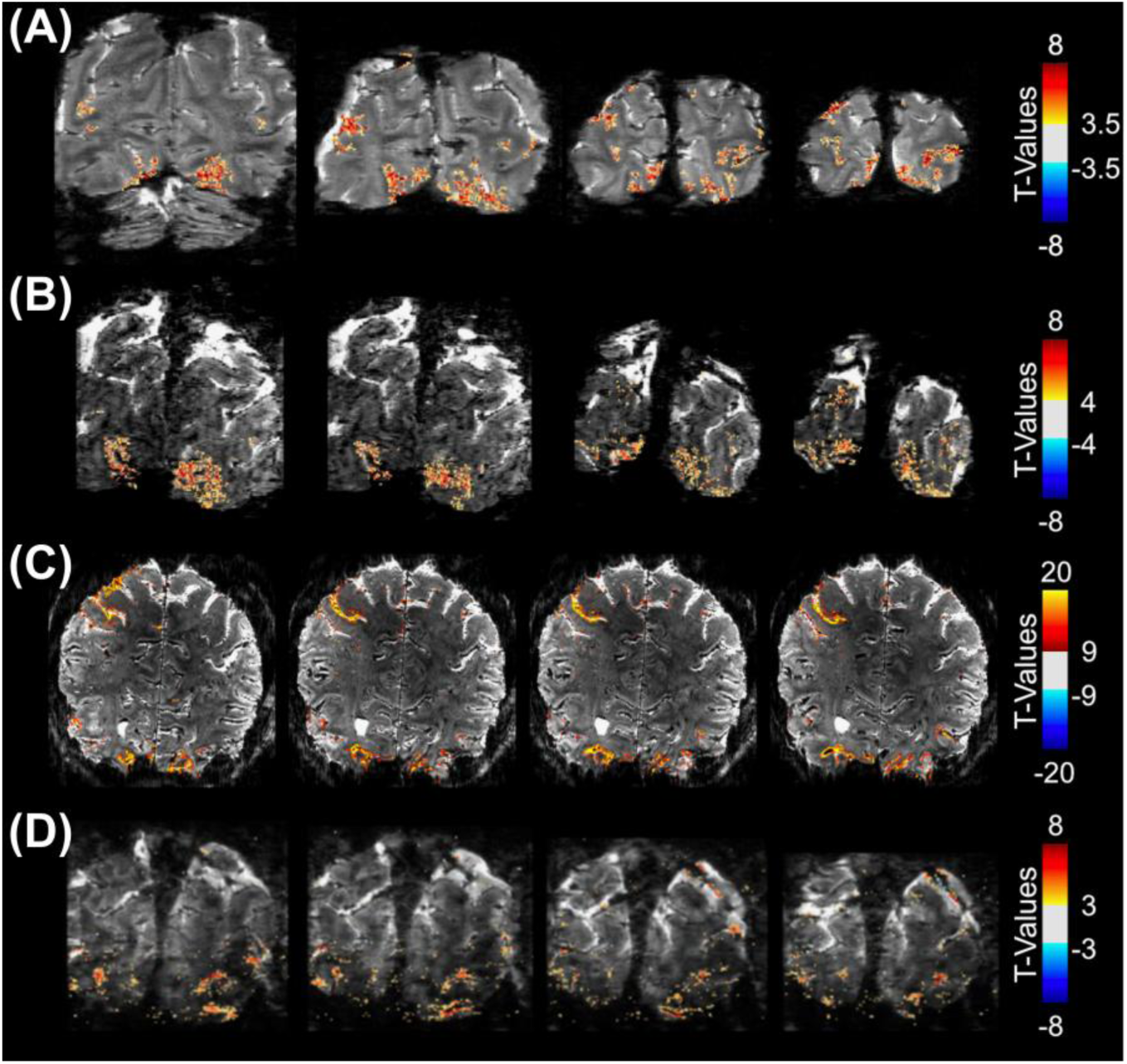
Functional MRI maps acquired during a visual stimulation task using four safety-validated head coils at 10.5T. Partial brain activation maps are shown for: (A) fMRI data acquired with the 8TxRx coil using a 2D GRE-EPI sequence at 0.54 × 0.54 × 0.8 mm³ resolution; (B) fMRI data acquired with the 16Tx/32Rx coil using a 2D GRE-EPI sequence at 0.4 × 0.4 × 0.6 mm³ resolution; (C) fMRI data acquired with the 16Tx/80Rx coil using a 3D GRE-EPI sequence at 0.5 mm isotropic resolution; and (D) fMRI data acquired with the 16Tx/128Rx coil using a 3D GRE-EPI sequence at 0.35 mm isotropic resolution.

#### 3.3.2 Diffusion MRI

Figure 10A–F presents dMRI metrics, including fractional anisotropy maps, primary diffusion orientation maps, and diffusion tensor estimations, acquired with 16Tx/80Rx and 16Tx/128Rx coils for the first time at 10.5T.

**Figure 10.**
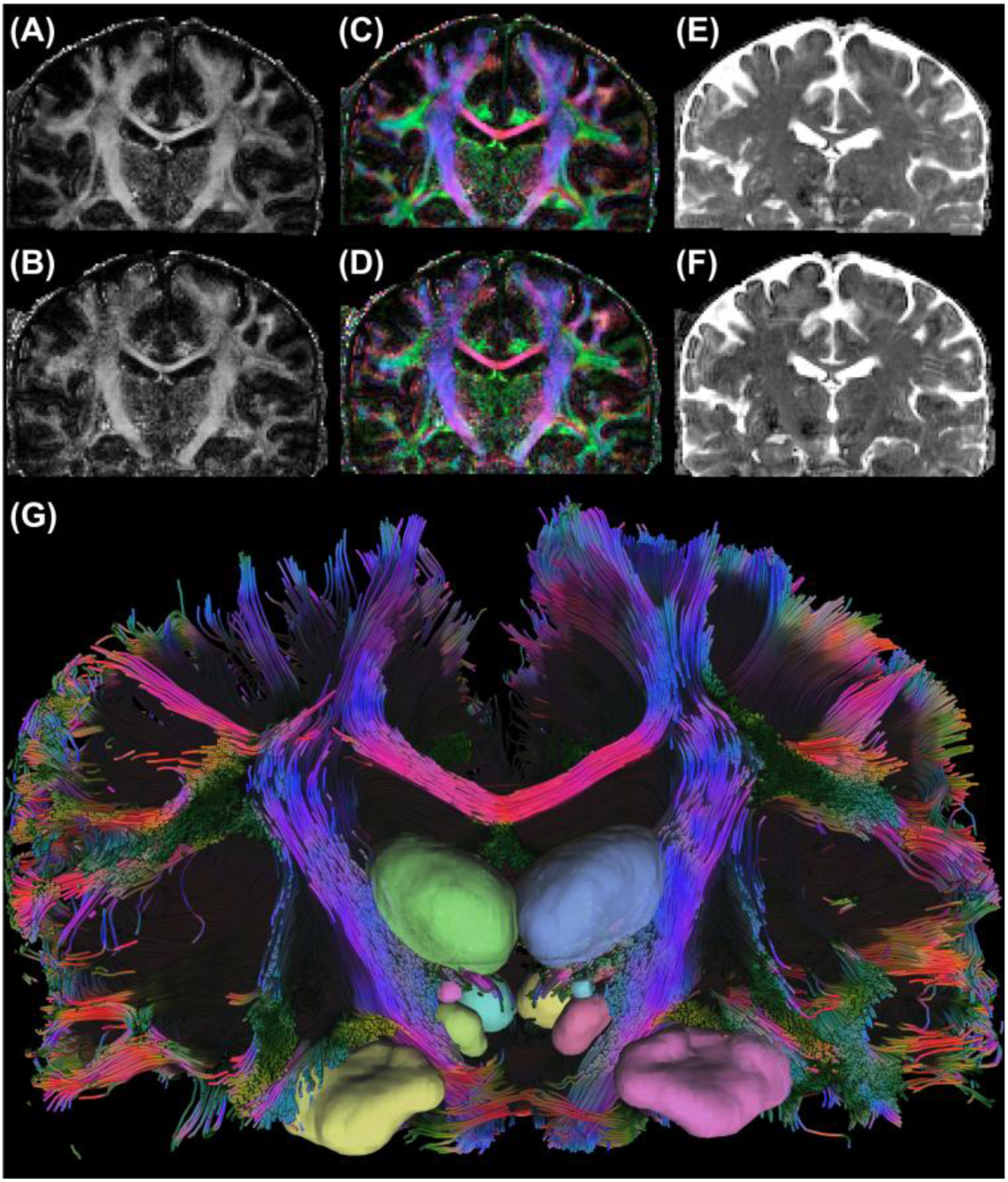
Diffusion MRI metrics following distortion correction using TOPUP and EDDY from FSL. (A–F): Top and bottom rows correspond to data acquired with the 16Tx/80Rx and 16Tx/128Rx coils, respectively. (A–B) Fractional anisotropy maps, (C–D) Primary diffusion orientation maps, and (E–F) Diffusion tensor estimations. (G) Tractography from the 16Tx/128Rx dMRI dataset, depicting major association, projection and commissural tracts, including the superior longitudinal fasciculus, corticospinal tract and corpus callosum.

Major association, projection, and commissural tracts are illustrated in Figure 10G, and additional details are presented in Supporting Information Figure S2.

## 4 DISCUSSION

In this study, a numerical technique was proposed as an alternative to the challenging MRT tests for quantifying EM modeling uncertainty in predicting *pSAR*_10*g*_ generated by custom-built multi-channel RF coils. The proposed technique was evaluated for performance using head and body multi-channel RF coils. It was also employed in the safety validation of three high-channel-count 10.5T head coils, which were subsequently used for in vivo functional and diffusion head imaging.

Specific discussions on the proposed technique and its implementation in this study are provided as follows:

### 4.1 Safety Validation of Multi-Channel RF Coils

#### 4.1.1 *e*_*EMM*_ Modeling Uncertainty Calculation

The proposed technique propagates the error between simulated and measured *B*^+^ to the error in simulated

*pSAR*_10*g*_ using Monte Carlo simulations. This technique is based on the hypothesis that per-channel excitations constitute the kernels of the excitation space. Building on this assumption, the *B*^+^ excitation space in the vicinity of a given mode of excitation can be generated by perturbing the complex weights of that mode. Quantifying the error between each perturbed mode and the original mode of interest provides insight into how that region of the excitation space is positioned with respect to the experimentally generated mode of interest. This concept forms the fundamental basis of the proposed technique. Since SAR and *B*^+^are correlated through their underlying electric and magnetic fields, governed by Maxwell’s equations, the *B*^+^error spaces can be translated into the SAR error spaces. Using the same rationale, the boundaries of the *B*^+^error region—which encompass all perturbed modes with *N*𝑅𝑀𝑆*E* values equal to or lower than the original *N*𝑅𝑀𝑆*E* between simulated and experimental *B*^+^maps—can be propagated to the SAR error region. The deviations of *pSAR*_10*g*_ of the perturbed modes within this region from the intended *pSAR*_10*g*_ of the mode of interest represent the uncertainty in the EM simulation when predicting *pSAR*_10*g*_.

In the proposed technique, the *pSAR*_10*g*_ error region encompasses all hypothetical excitation modes in the vicinity of the mode of interest that exhibit similar *B*^+^deviations from the simulated mode of interest as the experimentally acquired mode. Consequently, the *e*_*EMM*_ quantified using this approach represents the upper bound of the EM simulation error in predicting *pSAR*_10*g*_, making it a conservative yet safe estimate.

It is important to note that the proposed technique is not a replacement for MRT in applications requiring direct temperature change measurements in tissues or samples, such as hyperthermia treatments^58^ and RF ablation.^59^ Instead, it serves as an alternative approach to challenging, commonly used MRT-based tests for error quantification in EM models during the safety validation process of multi-channel RF coils.

#### 4.1.2 Validation Workflow

One of the primary parameters used to evaluate the agreement between simulation models and real-life RF coils is the S-matrix. While benchtop S-parameter measurements—as used in this study—provide reasonably accurate data, in-situ measurements inside the scanner bore could further improve EM modeling accuracy by accounting for complex in-bore components, such as the patient table and gradient shield. Previous studies at 7T have demonstrated that in-situ S-parameter measurements can significantly impact *pSAR*_10*g*_ predictions.^60^ However, implementing this approach presents challenges, primarily related to accessing and analyzing forward and reflected power data obtained from vendor-installed directional couplers at the output of the RF power amplifiers.

Another key factor in improving the agreement between EM simulations and experimental results is the incorporation of transmitter feed cable and T/R switch effects into the simulation/co-simulation process. These components primarily impact the measured and simulated *B*^+^ data in two ways:

***A. Signal Loss in the Transmit Chain:*** Feed cables and T/R switches introduce additional losses in the Tx chain, which can lead to significant discrepancies in the magnitude of simulated and measured *B*^+^ maps. For instance, in this study, the three 10.5T head coils exhibited a ∼1 dB loss.
1. ***B. Phase Differences Across Tx Channels:*** Variations in feed cable lengths and phase delays induced by T/R switches create relative phase differences between the Tx coil channels. If these phase variations are not accounted for in simulations, they can cause substantial errors in the *B*^+^field distribution.

To mitigate these discrepancies, both losses and phase delays were bench-measured for the three coils and incorporated into the co-simulation step.

There are several approaches to defining the cost functions for global and local optimizations in Step 3 of the validation workflow (Section 2.1.1). To reduce computational load and avoid excessive local minima, in this study, global optimization was performed only on diagonal elements of the S-matrix (***Cost 1***). In contrast, for the local optimization, diagonal elements along with relevant off-diagonal elements (with non-negligible measured magnitudes of > -20 dB) of the S-matrix were used with different weights (***Cost 2***).

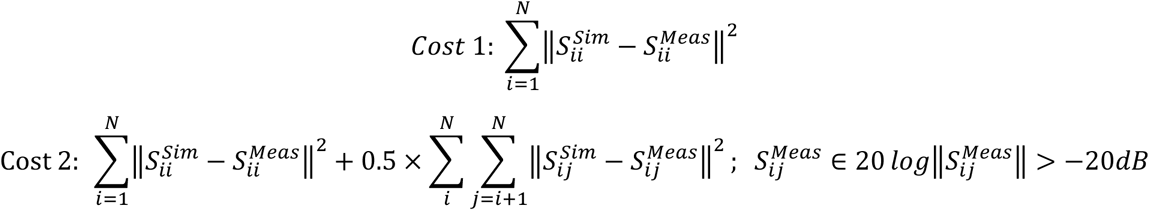

### 4.2 Performance Evaluation of the Proposed *e*_*EMM*_ Calculation Approach

#### 4.2.1 Reference to the MRT Technique

The performance of the proposed technique was evaluated using two excitation modes of a 16-channel transceiver body coil at 10.5T, which was previously safety-validated with *B*^+^mapping and MRT tests.^46^ The performance evaluation of the proposed technique was conducted using transceiver coils rather than pTx coils with dedicated receive-insert coils due to concerns about potential damage to sensitive receiver electronics from high-power MRT tests (typically ∼100W time-averaged power^46^). Furthermore, the performance evaluation tests using the body array coil included only two excitation modes to demonstrate the feasibility of estimating the upper bound of the *e*_*EMM*_, as discussed in Section 4.1.1 and shown in Figures 3 and 4. However, a comprehensive evaluation of the technique’s ability to estimate *e*_*EMM*_ across all modes would require testing a larger number of excitation modes, which is beyond the scope of this manuscript.

#### 4.2.2 Reference to the Temperature Probing Technique

In the performance evaluation of the proposed technique using the body array and MRT, the *e*_*EMM*_ for two independent excitation modes was calculated and validated (Steps 1–4 in Figure 1). However, their usability for other excitation modes was not investigated. To address this limitation, we *partially* verified the hypothesis that if a sufficiently large number of modes is included in the *e*_*EMM*_ calculation workflow (Step 5 in Figure 1), the final *e*_*EMM*_ would provide an adequate safety margin for any untested mode. This verification was performed using an 8-channel transceiver coil and 8 temperature probes, applying the full 5-step *e*_*EMM*_ calculation technique (Figure 1) across four different excitation modes. The highest *e*_*EMM*_ among the four modes was then applied to the 10g-averaged SAR values at the temperature probe locations for three additional random excitation modes. As shown in Figure 6, the *e*_*EMM*_ calculated from four modes provided a sufficient safety margin for the *pSAR*_10*g*_ of the three random modes, which were independently selected from the initial set of four. It is important to note that in this test, a full Q-matrix of the phantom was not utilized for *e*_*EMM*_ and *pSAR*_10*g*_ calculations. Instead, only 8 voxels, corresponding to the temperature probe locations, were used.

While this verification was successfully demonstrated for three modes, determining the universal *e*_*EMM*_ (i.e., safely covering the uncertainty for any excitation mode) for a coil requires a statistically justified sample size of excitation modes (***M*** in Figure 1). For instance, in the case of the 16Tx/32Rx coil, randomly sampling the excitation space with at least ***M*** = 37 would represent the full *B*^+^ error space within a 95% confidence interval, with a margin of error below 5%. To elaborate, the *N*𝑅𝑀𝑆*E* between simulated and experimental *B*^+^maps was computed for 10^6^ random excitation modes of the 16Tx/32Rx coil. These *N*𝑅𝑀𝑆*E* values were used to construct the *B*^+^ error space, yielding a mean (𝜇) of 54.5% and a standard deviation (𝜎) of 8.4%. The minimum sample size (𝑛) required to estimate the population mean with a 95% confidence level (𝑧 = 1.96) and a 5% margin of error (*e*) was calculated using the following formula^61^:

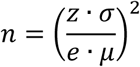

### 4.3 Case Studies: Safety Validation of three 10.5T Head Coils

In modeling the three 10.5T head coils, the gradient coil shield was excluded from the EM model, as prior simulations confirmed it had no detectable impact on the transmit fields.

Additionally, as highlighted in the Methods section, the high-channel-count receive-insert coils (32, 64, and 128 channels) were excluded from the final EM simulation models. This decision was justified through both numerical simulations and experimental measurements. For the numerical evaluation of this decision, the receive arrays’ loop conductors were included inside the transmit array coil, with all receiver loops terminated by high-input-impedance (∼3kΩ) to mimic the detuning circuit. The results showed negligible impact on the transmit fields, leading to the conclusion that the feed cables and receiver electronics are the primary contributors to Tx-Rx interactions, which are difficult to model in EM simulations. For the experimental evaluation of this decision, *B*^+^maps were acquired with and without the receive-insert in place, which again showed minimal impact.^53, 54^ It is noteworthy that, in the case of the 16Tx/128Rx coil, Tx-Rx interactions were further minimized by^54^: **1)** Implementing a network of current traps along the feed cables at ∼𝜆/16 intervals. **2)** Relocating electronics and minimizing their surface area to reduce interference.

As shown in Figure 8, the 16Tx/32Rx coil exhibited significantly higher EM modeling uncertainty compared to its higher- Rx-channel counterparts, despite having fewer receive channels—which might otherwise suggest a simpler model. This increased uncertainty is attributed to the modeling complexity of its 16Tx transmit array, which employed a transformer-based decoupling strategy between all neighboring elements. The elevated uncertainty is not related to the receive-insert array. In fact, the challenges associated with modeling transformer-decoupled loops served as a key motivation for transitioning to self-decoupled loop designs in the 16Tx/80Rx and 16Tx/128Rx coils.

As suggested in a previous study,^36^ intersubject variability (*e*_*ISV*_) can be either calculated individually for each coil or adopted from relevant literature. In this study, a 50% *e*_*ISV*_, based on 7T literature,^40^ was used to compute the safety factor for the three 10.5T head coils. However, recent studies,^42, 46^ including our own,^62^ indicate that *e*_*ISV*_ can vary significantly across different coils and field strengths. This suggests that a case-by-case evaluation of *e*_*ISV*_ would be a more reliable approach for safety factor determination.

## 5 CONCLUSIONS

In this study, we proposed a numerical technique for estimating EM modeling uncertainty (*e*_*EMM*_) in the prediction of *pSAR*_10*g*_ generated by multichannel RF coils, commonly used in UHF MRI applications. The effectiveness of this technique was evaluated through MRT tests and temperature probe measurements, both of which demonstrated that the proposed technique provides a conservative estimate of *e*_*EMM*_. While the technique resulted in some degree of overestimation, it successfully enabled the safety validation of three high-channel-count 10.5T head coils. Importantly, the conservatism in the derived safety factors did not preclude in vivo functional imaging of the human brain at an unprecedented 0.35 mm isotropic resolution, nor did it prevent acquisition of the first human diffusion MRI data at 10.5T.

## Acknowledgment

This research was funded by National Institutes of Health (NIH) grants: P41 EB027061, UM1 NS132207, R01 NS115180, R01 NS136490, and R01 EB029985.

