## Supporting Information for "A Numerical Alternative to MR Thermometry for Safety Validation of Multi-Channel RF Transmit Coils"

#### Functional MRI (fMRI)

##### Data Acquisition and Processing

fMRI data with submillimeter resolution were acquired at 10.5T (presented in Figure 9 of the manuscript) using four safety-validated head coils: 8TxRx[1], 16Tx/32Rx[2], 16Tx/80Rx[3], and 16Tx/128Rx[4, 5]. A standard 12 seconds (for the 2D acquisitions) and 24-second (for the 3D acquisitions) on/off visual block design was employed for all scans, featuring a central target and a surrounding counterphase flickering checkerboard. The differences in stimulations times were implemented to account for differences in volume acquisition time. The 0.5mm isotropic acquisition also included a concomitant motor task (closing and opening the right hand during blocks of visual stimulation). NORDIC[6] denoising was applied to the complex valued time-series, as described by Vizioli et al.[7]

- **8TxRx Coil**:

  2D GRE-EPI, resolution = 0.54 × 0.54 × 0.8 $mm^3$, 25 slices, TE/TR = 25/2000 ms, iPAT = 3, and 6/8 Partial Fourier.

- **16Tx/32Rx Coil**:

  2D GRE-EPI, resolution = 0.4 × 0.4 × 0.6 $mm^3$, 34 slices, TE/TR = 25/2576 ms, iPAT = 4, and 5/8 Partial Fourier.

- **16Tx/80Rx Coil**:

  3D GRE-EPI, 0.5 mm isotropic resolution, 22 slices, TE/TR = 22/104 ms, VAT = 2500 ms, iPAT = 4, and 5/8 Partial Fourier.

- **16Tx/128Rx Coil**:

  3D GRE-EPI, 0.35 mm isotropic resolution, 40 slices, TE/TR = 22.6/104 ms, VAT = 5016 ms, iPAT = 4, and 5/8 Partial Fourier.

All functional data preprocessing was performed using BrainVoyager, with procedures kept minimal and consistent across reconstructions. Specifically, slice scan timing correction was applied only to the 2D datasets using temporal sinc interpolation. 3D rigid-body motion correction was performed using spatial sinc interpolation, with all volumes from all runs aligned to the first volume of the first acquired run. Low-frequency drift removal was carried out using a general linear model (GLM) approach, employing a design matrix that included up to the third-order discrete cosine transform basis set. No spatial or temporal smoothing was applied.

Functional data were aligned to anatomical scans through manual adjustments and iterative optimization. Standard GLM analyses were then used to estimate percent signal change amplitudes and corresponding t-values elicited by the target and surround conditions (see Vizioli et al.[8] for more details).

##### RF Limitations

In general, SAR efficiency (i.e., defined as $\frac{B_1^+}{\sqrt{pSAR_{10g}}}$) decreases at higher field strengths,[9-11] and applying additional safety factors or overestimating $pSAR_{10g}$ aggravates this issue. One consequence was that our fMRI studies experienced up to ~40% underflipping, leading to suboptimal SNR. Several factors contribute to this problem:

***A. Conservative $e_{EMM}$ Estimation:*** The $e_{EMM}$s for the three 10.5T head coils were calculated in the range of ~40% to ~130%, resulting in conservative safety factors of ~2. Such high $e_{EMM}$s, which exceeded the ground truth $e_{EMM}$ in the evaluated scenarios (see Figures 3 and 4), stems from using the 99.9th percentile of the $pSAR_{10g}$ error region for $e_{EMM}$ quantification. Lowering this threshold could reduce safety factors, but further investigations with multiple excitation modes are required before making such a modification. Additionally, enhancing EM modeling accuracy could reduce the

$B_1^+$ $NRMSE$, and therefore, the $e_{EMM}$, which may be achieved through in-bore S-parameter measurements instead of bench-top measurements.[12]

***B. Intersubject Variability:*** The 50% $e_{ISV}$ applied to the three 10.5T head coils significantly contributed to the high safety factors (~2). This variability could be reduced using individualized models[13] or subject-specific deep learning-based SAR estimation techniques.[13, 14]

***C. $pSAR_{10g}$ vs. Temperature Limits:*** According to IEC guidelines,[15] the local SAR serves as a proxy for temperature increases which can cause tissue damage. However, SAR simulations do not account for the significant impact of perfusion on regulating body temperature, leading to an overestimation of the risks for tissue damage. This overestimation suggests that SAR limits should ultimately be replaced by temperature-based metrics.[16] Two potential approaches include: 1) Temperature matrices simulations incorporating perfusion effects,[17] instead of $Q$-matrices and 2) In vivo MRT techniques.[18]

***D. Overestimation in VOPs:*** VOP compression[19] is essential for real-time $pSAR_{10g}$ monitoring in pTx coils, but it also introduces a potential overestimation of pSAR. To comply with vendor-imposed "coil-file" size limitations, a ~10% overestimation (on $Q$-matrix eigenvalues) was applied in VOP compression in this study. However, in certain modes—such as CP mode in the three 10.5T head coils—this led to up to ~30% overestimation. To address this issue, modified VOP compression techniques have been introduced.[20]

***E. Parallel Transmission (pTx) Pulses:*** In our fMRI studies, static RF shimming was used with a minimum excitation inhomogeneity target and no $pSAR_{10g}$ constraints. More effective alternatives include $pSAR_{10g}$-constrained shimming[21] and dynamic shimming techniques, such as multi-spoke RF pulses, which have demonstrated superior performance by providing more degrees of freedom for pTx optimization.[22, 23]

#### Diffusion MRI (dMRI)

The first in vivo human brain dMRI data at 10.5T were acquired (presented in Figure 10 of the manuscript) using two safety-validated high-channel-count head coils: 16Tx/80Rx[3] and 16Tx/128Rx.[4] Whole-brain dMRI was performed using a 2D SE-EPI sequence with TE/TR = 63/10,000 ms, 1.05 mm isotropic resolution, iPAT = 4, matrix size = 200 × 200, and Receiver Bandwidth = 277600 Hz. The diffusion acquisition employed a two-shell q-space sampling scheme with b-values of 900 and 1800 s/mm², covering a total of 71 diffusion directions per shell.

Imaging was performed using an RF shimming solution, optimized by minimizing the coefficient of variation ($CoV$) of $B_1^+$ across the whole brain as a measure of transmit field inhomogeneity, while constraining $pSAR_{10g}$ for safety compliance:

$$\begin{array}{cc} min & CoV \\ subject\ to & pSAR_{10g} \leq target\ pSAR_{10g} \end{array}$$

where, target $pSAR_{10g}$ value was varied as a fraction of that for the CP mode. This shimming strategy produced the commonly-used $pSAR_{10g} - COV$ L-curve, which represents the trade-off between excitation homogeneity and safety limits (see Supporting Information Figure S1).

Data were collected with reversed phase-encode blips, producing image pairs with opposite distortion patterns. These were used to estimate the susceptibility-induced off-resonance field using a method similar to that of Andersson et al.,[24] implemented via TOPUP in FSL,[25] after which the two images were combined into a single distortion-corrected volume. Eddy current and motion-induced distortions, as well as outliers, were corrected using the EDDY tool, following the methodology described in the literature.[26, 27]

The diffusion data were reconstructed using generalized q-sampling imaging[28] with a diffusion sampling length ratio of 1.25. Deterministic fiber tracking[29] was performed, with augmented tracking strategies[30] implemented to enhance reproducibility. The anisotropy threshold was randomly selected between 0.5- and 0.7-times Otsu threshold. The analysis was conducted using DSI Studio (Hou, http://dsi-studio.labsolver.org). Major association tracts derived from tractography of the 16Tx/128Rx dMRI data are shown in Supporting Information Figure S2.

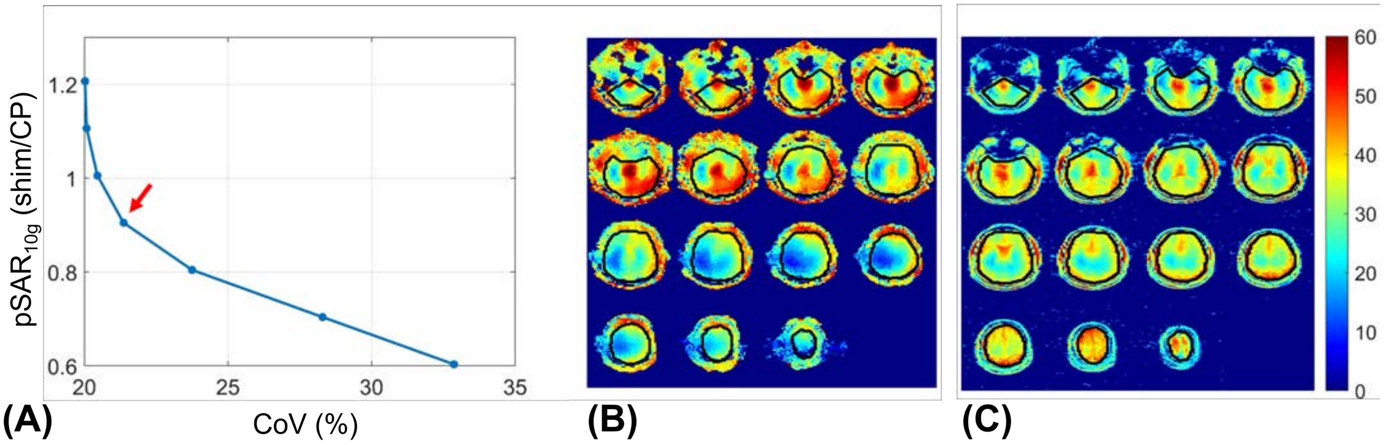


**Supporting Information Figure S1.** Summary of $pSAR_{10g}$-constrained excitation homogeneity RF shimming used for diffusion MRI at 10.5T with the 16Tx/128Rx head coil. **(A)** $pSAR_{10g} - COV$ L-curve resulting from iterative optimization, where the target $pSAR_{10g}$ was varied as a fraction of the value for the CP mode. The red arrow indicates the optimum RF shim solution selected for imaging. **(B–C)** Flip angle maps acquired using the AFI technique for **(B)** the CP mode and **(C)** the optimized shim solution corresponding to the red arrow in panel (A). The region of interest used for CoV calculation is outlined in black.

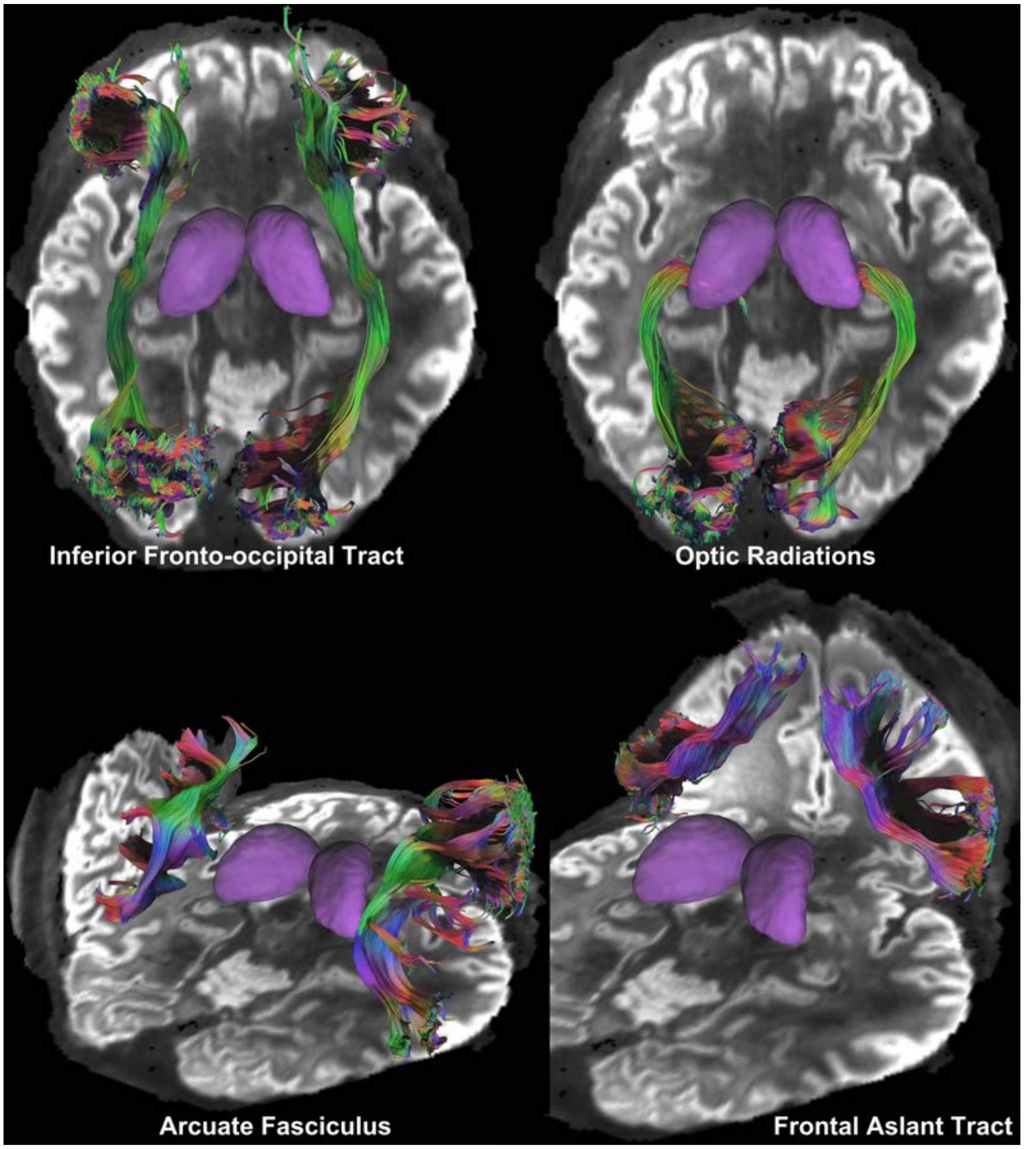


**Supporting Information Figure S2.** Tractography from the 16Tx/128Rx dMRI data depicting major association tracts. The diffusion data were reconstructed using generalized q-sampling imaging with a diffusion sampling length ratio of 1.25. A deterministic fiber tracking algorithm was used with augmented tracking strategies to improve reproducibility. Autotrack was used to automatically identify tracts with a distance tolerance of 24.00 (mm) in the ICBM152 space by comparing trajectories with a tractography atlas. Topology-informed pruning was applied to the tractography with 8 iterations to remove false connections. The anisotropy threshold was randomly selected between 0.5 and 0.7 Otsu threshold. The analysis was conducted using DSI Studio (Hou, http://dsi-studio.labsolver.org).

### BIBLIOGRAPHY to SUPPORTING INFORMATION

**14.** Kim J, Sadeghi-Tarakameh A, Torrado-Carvajal A, Eryaman Y. A Novel Specific Absorption Rate Prediction Framework Using Multi-Task Feedback Generative Adversarial Learning: Application to 10.5 T Head MRI. Paper presented at: Proc Int Soc Mag Reson Med. ; 2021, 2021; In: Proc Int Soc Mag Reson Med. Online.

**30.** Yeh FC. Shape analysis of the human association pathways. *Neuroimage.* Dec 2020;223.
